# Synaptotoxic forms of amyloid-β and α-synuclein act through a common pathway

**DOI:** 10.1101/2025.02.13.638131

**Authors:** E. Ribe, A. Ghosh, W. Paslawski, K. Savory, C. Ballard, P. Morin, D. Cash, F. Hirth, G. Williams, P. Svenningsson, D. Aarsland, R. C. M. Siow, R. Killick

**Affiliations:** Centre for Healthy Brain Aging, King’s College London, Institute of Psychiatry, Psychology & Neuroscience, London, SE5 9RT, UK; School of Cardiovascular and Metabolic Medicine and Sciences, King’s BHF Centre of Research Excellence, Faculty of Life Sciences and Medicine, King’s College London, 150 Stamford Street, London SE1 9NH, U.K; Department of Clinical Neuroscience, Neuro Svenningsson, Karolinska Institute, 171 76 Stockholm, Sweden; College of Medicine and Health, University of Exeter, Exeter EX1 2UL, UK; Department of Neurology, Boston University Chobanian & Avedisian School of Medicine, Boston, MA, USA; BRAIN Centre (Biomarker Research and Imaging for Neuroscience), Department of Neuroimaging, King’s College London, The James Black Centre, 125 Coldharbour Lane, London SE5 9NU, UK; Department of Basic & Clinical Neuroscience, King’s College London, Institute of Psychiatry, Psychology & Neuroscience, Maurice Wohl Clinical Neuroscience Institute, London SE5 9RX, UK; Wolfson SPaRC, The Wolfson Wing, Hodgkin Building, King’s College London, London SE1 1UL, UK; Department of Physiology, Anatomy and Genetics, Medical Sciences Division, University of Oxford

**Keywords:** Synuclein, amyloid, dendritic spine, synapse, Parkinson’s, Alzheimer’s, Wnt, planar cell polarity, Daam1, Rock, fasudil

## Abstract

It has been suggested that α-synuclein (αSyn), a major player in Parkinson’s disease (PD), plays a role in Alzheimer’s disease (AD). Several reports have also concluded that αSyn and amyloid-β (Aβ) are mechanistically linked, although how is unclear. Synapse loss is an early feature in both PD and AD and held to be the driver of both diseases. We have previously uncovered a signalling pathway required for Aβ-driven dendritic spine loss - a non-canonical branch of Wnt signalling known as the Wnt/Planar Cell Polarity (Wnt/PCP) pathway. We asked if a synaptotoxic form of αSyn known to impact dendritic spines, the A53T autosomal dominant PD mutant form of αSyn (A53T-αSyn), might act on synapses through the same pathway. Here, by blocking all Wnt activity with the porcupine inhibitor IWP2, we show that A53T-αSyn-driven spine loss is Wnt-dependent. By silencing *Daam1*, which is unique to Wnt/PCP, we show that A53T-αSyn spine loss is *Daam1*-dependent. Finally, using the pan-ROCK inhibitor fasudil indicates the mechanism also involves ROCK1/2, which *Daam1* signals to via RhoA to modulate actin cytoskeletal dynamics within dendritic spines. Together, these observations indicate that A53T-αSyn-driven spine loss involves the Wnt/PCP pathway, the same pathway that mediates Aβ synaptotoxicity. This indicates that Aβ and αSyn are mechanistically connected and that a common pathway is responsible for synapse loss in AD and PD. It also begins to explain why this group of neurodegenerative diseases have many features in common and suggests that drugs which target Wnt/PCP could be of benefit for both AD and PD.

## Introduction

Amyloid-β (Aβ) is central to Alzheimer’s disease (AD). As insoluble aggregates, Aβ contributes to senile plaques in the brain, which along with tau pathology in the form of neurofibrillary tangles are the two defining neuropathological hallmarks of AD. It is however the soluble oligomeric forms of Aβ that exert toxic effects upon synapses leading to synapse loss, widely held to be an early event in the underlying disease process driving AD(1).

The characteristic hallmark of Parkinson’s disease (PD) and the related synucleinopathies, PD with dementia (PDD) and dementia with Lewy bodies (DLB), are Lewy bodies and Lewy neurites, intraneuronal deposits containing highly aggregated forms of the α-synuclein (αSyn) protein. It is now apparent that Lewy pathology is also present in a majority of AD cases(2, 3), and when predominant in the amygdala, correlates with advanced hippocampal neuronal loss(4). Furthermore, increased levels of soluble αSyn in cerebrospinal fluid (CSF) are associated with faster rates of cognitive decline in AD(5, 6), and positivity in the seed amplification assay (SAA) for misfolded αSyn, a pathology-specific marker for Lewy body disease (LBD)(7), is associated with increased brain Aβ levels as measured by PET in mild cognitive impairment and AD(8). Conversely, a recent histopathological study found Aβ pathology in 75% of PD cases, with higher levels being associated with accelerated disease progression(9), and in a separate study CSF-based AD biomarkers predicted levels of cognitive impairment in PD(10).

A considerable number of reports have also suggested that the αSyn protein and the Aβ peptide are in some way functionally connected(11–17), although how has not yet been determined. These and many other observations have led some to suggest that αSyn may directly contribute to the neuropathogenesis of AD(18), while they equally suggest Aβ may play an important role in Lewy body diseases (LBDs).

The effects of Aβ on dendritic spines in rodent primary neuronal cultures have been studied extensively(19–22). Dendritic spines are small protrusions found along dendrites(23), constituting the postsynaptic element of excitatory glutamatergic synapses(24). They play a crucial role in synaptic plasticity and are considered part of the biological basis of learning and memory(24–26). A failure in the proper regulation of actin cytoskeletal dynamics within dendritic spines has been suggested to play a major role in the loss of synaptic connectivity seen in AD(25).

Wnt signalling plays a major role in synapse formation, growth, stabilisation and retraction(27–30). While a complete understanding of the mechanisms underlying Aβ-driven synapse loss has yet to be obtained, we(31, 32), and others(33), have shown that the synaptotoxic effects of Aβ are dependent upon, and mediated through, Wnt pathway component Dickkopf-1 (Dkk1). In this context Dkk1 acts as an antagonist of the canonical Wnt/β-catenin pathway by binding and removing LRP6 from the Frizzled co-receptor complex(34), its presence in which is needed for canonical Wnt to proceed. Importantly, this has a second effect, that of allowing activity in the non-canonical Wnt pathways to proceed, which the LRP6 interaction with Frizzled holds in check(35, 36). In this way, Dkk1 binding LRP6 results in the concomitant activation of the Wnt/Planar Cell Polarity (PCP) pathway, the main non-canonical branch of Wnt signalling involved in the regulation of actin cytoskeletal dynamics(37–39). Our previous work has shown that Aβ, via Dkk1, activates Wnt/PCP and that it is this aberrant modulation of Wnt/PCP which mediates the synaptic effects of Aβ on dendritic spines, leading to spine retraction and synapse loss(31, 32).

Synapse loss is also an early event in PD, and as in AD involves a dysregulation of glutamatergic receptors(40). While the αSyn protein is abundantly expressed at presynaptic terminals and its role in neurotransmitter release studied intensively(41), it is well documented that reductions in dendritic spine density occur in the striatal medium spiny neurons in rodent models of PD(42) and in post-mortem PD brain(43, 44). Golgi staining has also revealed an almost complete loss of dendritic spines adjacent to very small presynaptic deposits of αSyn in frontal cortex in DLB(45). Furthermore, in independent studies of transgenic mouse models carrying the autosomal dominant A53T familial PD mutation in *SNCA*, the human A53T αSyn protein was found to drive postsynaptic impairment(46), have deleterious effects on dendritic spines(47) and ‘support the role of postsynaptic degeneration as an early feature in synucleinopathies’(48).

Here, we investigated whether the effects of A53T-αSyn on dendritic spines might involve the Wnt/PCP pathway. We used an oligomerised recombinant human A53T-αSyn protein, adding it directly to the growth media of rat primary cortical cultures to determine its effects on dendritic spines. Our results provide evidence that A53T-αSyn triggered dendritic spine loss is mediated via the Wnt/PCP pathway, the same pathway that mediates Aβ synaptotoxicity, suggesting they are indeed functionally related, acting in a common pathogenic pathway that may underpin synapse loss in AD and PD.

## Methods

### Cortical cultures

Mixed, neuronally enriched, E18 rat primary cortical cultures were prepared from Sprague-Dawley E18 rat embryo brains. Brains were collected in HBSS on ice, meninges were removed, then cortices dissected and dissociated in 2.5% trypsin (Invitrogen 15090046) for 30 min. DNase 1 (Sigma D5025) was added and cells were allowed to settle before removing supernatant and triturating in HBSS with Albumax (Invitrogen 11020-013), trypsin inhibitor (Sigma T9003) and DNase1. Cell suspensions were then diluted in Neurobasal media (25030024) + B27 supplement (17504044) + GlutaMax (35050–038) + penicillin/streptomycin (P4333) - all obtained from Thermo Fisher -, counted, and seeded into 12 well plates containing 18 mm 1.5H precision coverslips (0117580 Marienfeld), coated with poly-D-lysine (P7886 Merck life Sciences), at 0.1 mg/mL in water, at a density of 300,000/well. After 4 days in vitro (d.i.v.), 50% percent media changes were performed twice weekly for 1 week, followed by 25% media changes twice weekly until the desired time in culture was reached (23-24 d.i.v.).

### eGFP transfection

Cultures were transfected with an expression construct encoding enhanced green fluorescent protein (eGFP) driven by the synapsin-1 promoter using Lipofectamine 2000 resulting in the desired low transfection efficacy (<0.01%). The exogenous expression of eGFP in just 15-20 neurons per cover slip allows for the imaging of individual dendrites and dendritic spines without the need for further labelling.

### Preparation of A53T-αSyn recombinant protein

A53T-αSyn monomers were prepared as described previously, with minor modifications(49). Briefly, human A53T-αSyn in pET11-D was expressed in *E. coli* BL21(DE3) competent cells using an auto-induction method. Cells were harvested by centrifugation and resuspended in osmotic shock buffer (20 mM Tris-HCl, pH-7.2, 40% sucrose), incubated for 10 min and centrifuged again. The pellet was re-suspended in ice-cold deionised water, with subsequent addition of saturated MgCl_2_, and briefly incubated on ice. The periplasmic fraction of the cell lysate was collected, and the majority of unwanted proteins precipitated by adjusting the pH to 3.5 with 1 M HCl. Soluble proteins were collected by centrifugation and the pH of the obtained supernatant was adjusted to pH-7.5 with 1 M NaOH. The solution was filtered and fractionated on a Q-Sepharose column connected to an ÄKTA Go system (GE Healthcare) using a rising concentration of NaCl. Fractions containing A53T-αSyn were identified by SDS-PAGE and pooled. Further, high molecular weight aggregates were removed by filtration through a 30 kDa cut-off filter, to obtain only monomeric forms of αSyn, and analysed with SDS-PAGE to ensure purity. Next, to remove endotoxins, the solution was incubated with Lipid Removal Agent (Merck) according to manufacturer’s instructions and finally dialysed against deionised water. The A53T-αSyn concentration was determined by BCA assay (Thermo Fischer Scientific), protein was aliquoted, lyophilised and stored at −20°C.

### Oligomerisation of A53T-αSyn

To generate A53T-αSyn oligomers the recombinant protein was resuspended in PBS at 120 µM, aliquoted and stored at −80°C. Following a modified protocol of Bate et al.(12), who previously reported that the synaptic effects of Aβ and αSyn are synergistic, on the day prior to use an aliquot was thawed and allowed to aggregate at 37°C for 24 h. After chilling on ice, the solution was sonicated for 10 min and then diluted directly into culture media at 1 in 40 to achieve a final working concentration of 3 µM. A western blot showing the oligomerised A53T-αSyn prep run under non-denaturing conditions is shown in figure 1A.

**Figure 1.**
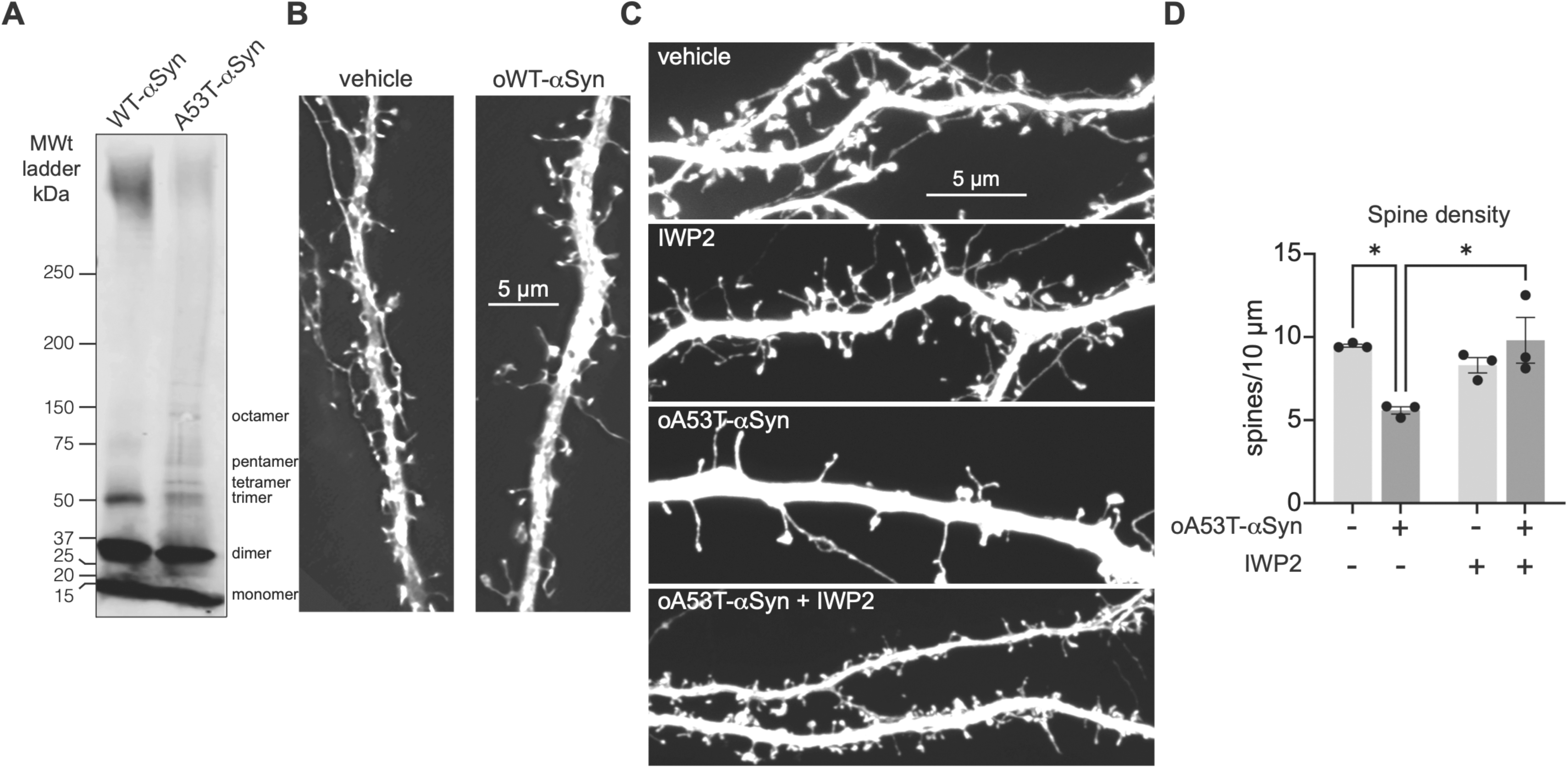
Oligomeric A53T-αSyn-driven dendritic spine loss is Wnt-dependent. **A,** western blot of oligomerised WT and A53T αSyn recombinant proteins showing potential oligomeric forms. **B**, E18 rat cortical cultures transfected with synapsin-driven eGFP plasmid after 24 days in vitro (d.i.v.) and next day treated with vehicle or 3 µM oligomerised WT-αSyn recombinant protein for 24 hours, fixed and imaged by super resolution microscopy. **C,** corresponding cultures as in B were treated overnight with 1 µM IWP2 and next day treated with 3 µM oA53T-αSyn recombinant protein for 24 hours, processed and imaged as in B. Images show representative levels of dendritic spine density across the four conditions. **D**, quantification of dendritic spine density indicates that oA53T-αSyn drives substantial dendritic spine loss, which is significantly reduced by blocking all Wnt signalling activity with IWP2. Error bars indicate SEM *=p<0.05

### Preparation of inhibitors

Fasudil HCl (Selleck) was solubilised in water at 2 mM and added to cultures at a final concentration of 10 µM 15 min prior to adding A53T-αSyn. IWP2 was solubilised in DMSO at 10 mM and used at 1 µM overnight before the addition of A53T-αSyn.

### *Daam1/2* siRNA knockdown

Pen1-coupled small interfering RNA (siRNA) oligonucleotides to rat *Daam1* and *Daam2* were generated and characterised previously(31). In brief, siRNA sequences were designed according to the protocol of Reynolds et al., 2004(50) with the inclusion of dTdT 3° overhangs and a 5°-thiol modification to the sense strand, then coupled to the cell penetrating peptide, Pen1, as described in Davidson et al., 2004(51), in order to obtain very high transfection efficiency of all cell types present in the primary cortical cultures. *Daam2* is not detectable in rat cortical cultures as we previously reported(31), as such the *Daam2* siRNA was used here as a control for the *Daam1* siRNA.

### Imaging

Transfected and fixed cultures were imaged using a Nikon inverted super resolution instant structured illumination (iSIM) microscope with a Hamamatsu flash 4.0 sCMOS camera and 100X (Numerical Aperture 1.35) silicone immersion lens housed in the Wohl Cellular Imaging Centre. Dendritic spines were quantified using the Dendritic Spine Counter plug-in, in ImageJ/FIJI.

### Statistical analysis

For each condition, 30-40 dendrites from at least three separate experiments were used for statistical analysis, with averages from a single rat prep considered as a single biological replicate. Experiments were carried out blind to condition. Data was analysed in GraphPad Prism, using unpaired t-tests or one-way ANOVA with post hoc Tukey’s multiple comparisons testing. Normality of data was assessed using Shapiro-Wilk test. Analyses having P<0.05 or lower were considered biologically significant.

## Results

To investigate if A53T-αSyn might impact dendritic spines via Wnt/PCP we employed the same model we previously used to investigate Aβ synaptotoxicity(31, 32), E18 rat cortical cultures maintained for 25-26 days *in vitro* (d.i.v.). At this time functional glutamatergic synapses have formed, and dendritic spines, the post synaptic element of such synapses, are detectable. To allow the visualisation and quantification of individual dendritic spines by super resolution iSIM microscopy, cultures were typically transfected after 23-24 d.i.v. with a synapsin-1 promoter-driven eGFP expression plasmid. To enhance the synaptotoxicity of the A53T-αSyn recombinant protein it was oligomerised (see methods) as shown by western blot (figure 1A). A dose and exposure time were then empirically determined, finding that exposure to 3 µM oligomeric A53T-αSyn (oA53T-αSyn) for 24 h resulted in ∼50% spine loss. At this dose and exposure time oligomerised WT-αSyn protein had no discernible effects on spine density (figure 1B).

To determine if the effect of oA53T-αSyn on dendritic spine loss involves Wnt signalling we employed IWP2, a small molecule inhibitor of the enzyme protein-serine O-palmitoleoyltransferase, the mammalian homologue of *Drosophila* Porcupine(52). Porcupine is responsible for the palmitoylation of all Wnt ligands, a step in Wnt protein maturation necessary for biological activity(53). Cultures were transfected with an eGFP plasmid, treated the next day with 1 µM IWP2 overnight and on the following day exposed to 3 µM oA53T-αSyn for 24 h. As shown in figure 1C & 1D, in the presence of IWP2 the effects of oA53T-αSyn on dendritic spine density were significantly reduced, indicating that the mechanism through which oA53T-αSyn-drives dendritic spine loss is dependent on Wnt signalling.

Next, to determine if oA53T-αSyn impacts dendritic spines specifically via the Wnt/PCP pathway we silenced *Daam1* and *Daam2*, key elements of the Wnt/PCP pathway acting below Dishevelled for which no function outside the Wnt/PCP pathway has, to the best of our knowledge, yet been reported. In rat cortical cultures only *Daam1* is expressed(31). As such we used si*Daam2* as a control for si*Daam1*. Cultures were transfected with the eGFP expression plasmid, and next day treated with si*Daam1* or si*Daam2* coupled to Pen-1, allowing highly efficient silencing across all cells in the culture(51). On the following day cultures were exposed to 3 µM oA53T-αSyn for 24 h. Imaging and spine quantification revealed that si*Daam2* did not impact oA53T-asyn driven spine loss, however, silencing *Daam1* potently and significantly reduced oA53T-αSyn-driven spine loss (figure 2). This indicates that the effects of oA53T-αSyn on dendritic spines involves *Daam1* and activation of the Wnt/PCP pathway. *Daam1* relays the Wnt/PCP signal via RhoA to ROCK, which then modulates actin cytoskeletal dynamics to regulate dendritic spine morphology(54).

**Figure 2.**
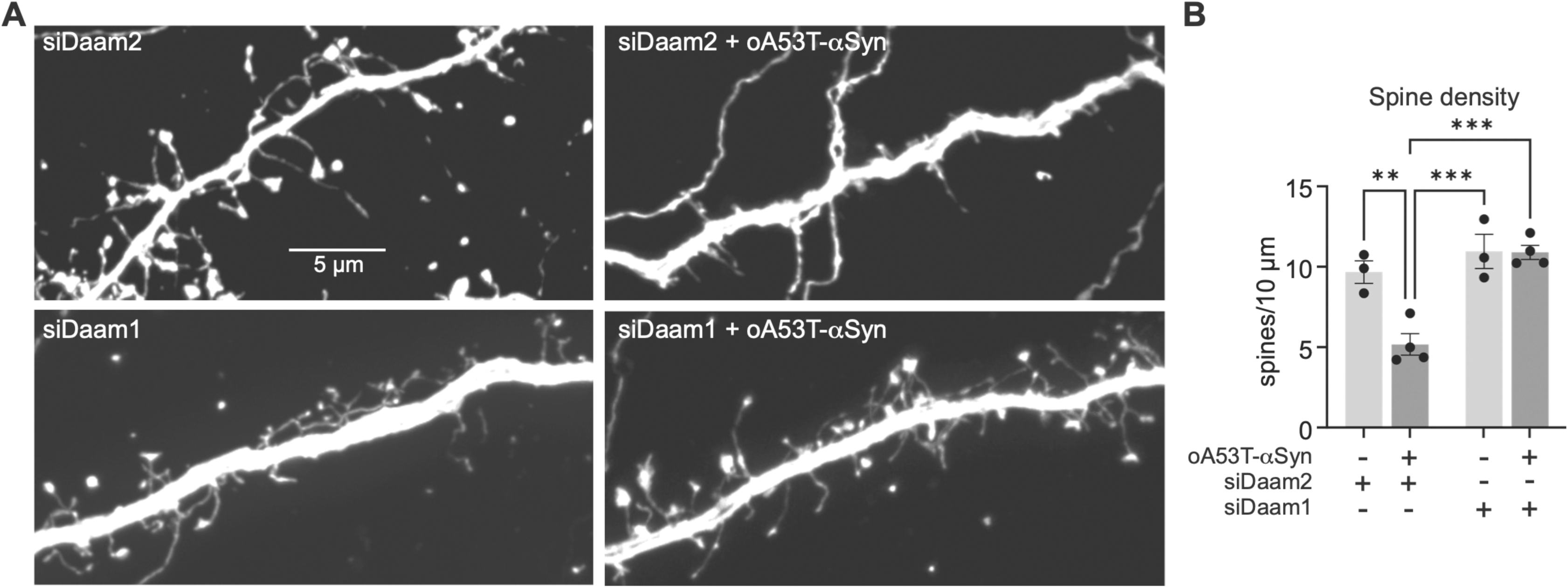
oA53T-αSyn-driven dendritic spine loss is Daam1-dependent. **A,** E18 rat cortical cultures as in figure 1B, above, were treated with Pen-1 coupled *Daam1* or *Daam 2* siRNA at 25 d.i.c. and next day treated with 3 µM oA53T-αSyn recombinant protein for 24 hours, fixed and imaged. Images show representative levels of dendritic spine density across the four conditions. **B**, quantification of dendritic spine density indicates that oA53T-αSyn-driven dendritic spine loss is *Daam1*-dependent. Error bars indicate SEM **=p<0.01, ***=p<0.001

To determine if oA53T-αSyn drives spine loss via ROCK we pre-treated cultures with the pan-ROCK inhibitor fasudil at 10 µM, or with vehicle, for 15 min prior to the addition of 3 µM oA53T-αSyn to the culture media for 24 h. As shown in figure 3A & 3B, fasudil blocked oA53T-αSyn-driven spine loss, as it also does Aβo-driven spine loss(31).

**Figure 3.**
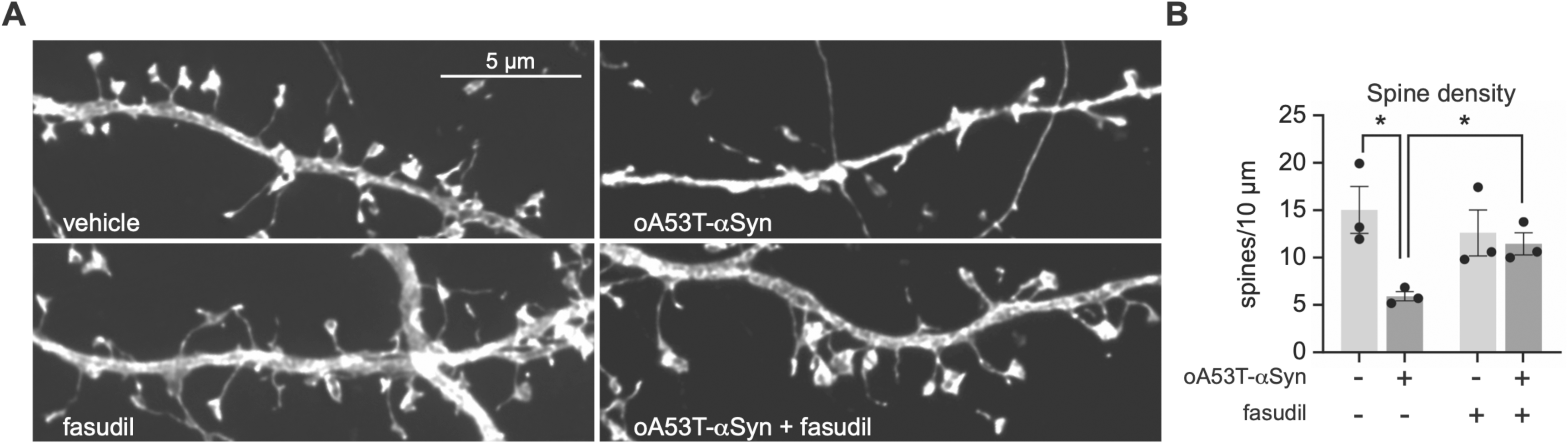
oA53T-αSyn-driven dendritic spine loss involves ROCK1/2. **A,** E18 rat cortical cultures were treated at 26 d.i.c. with 10 µM fasudil or vehicle for 15 min prior to exposure to 3 µM oA53T-αSyn recombinant protein for 24 hours and imaged, as above. Images show representative levels of dendritic spine density across the four conditions. **B**, quantification of dendritic spine density indicates that oA53T-αSyn-driven dendritic spine loss involves ROCK1 and or ROCK2 activity. Error bars indicate SEM *=p<0.05

## Discussion

Our findings demonstrate that in rat primary cortical neurons oligomeric A53T-αSyn drives the loss of dendritic spines in a Wnt-, Daam1-, and ROCK1/2-dependent manner, together pointing to an involvement of the Wnt/PCP pathway. As a similar function has been reported for oligomeric Aβ_1-42(31)_, this indicates that a form of αSyn which causes early onset familial PD, acts on synapses via the same pathway as Aβ.

Accumulation and aggregation of oligomeric Aβ_1-42_ is considered the causative agent of both familial and sporadic forms of AD, whereas accumulation and aggregation of oligomeric αSyn is considered the causative agent of both familial and sporadic forms of PD, and likely PDD and DLB neuropathologies too(45). The observed Wnt/PCP-mediated synapse loss triggered by accumulating oligomeric Aβ_1-42(31)_ or oligomeric A53T-aSyn (this report) thus indicates that these proteinopathies, which together account for a majority of cases of neurodegenerative disease, have a common underlying synaptopathic mechanism (figure 4).

**Figure 4.**
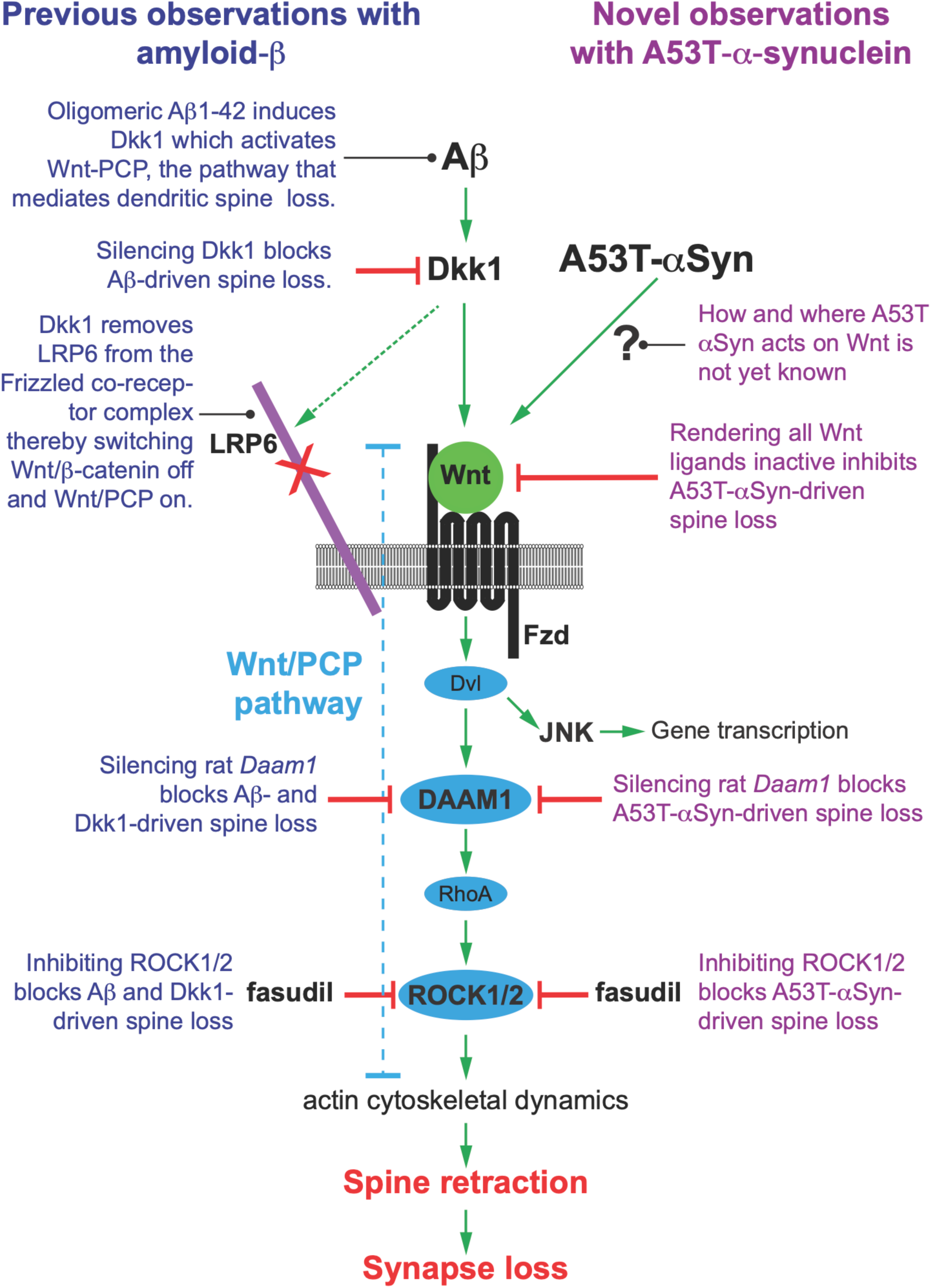
Schematic of common αSyn-Aβ pathway. Summary of findings and current hypothesis showing previous observations concerning the effects of Aβ triggered synapse loss (left, in blue font) and reported novel observation of oA53T-αSyn triggered synapse loss (right, in purple font), both of which are mediated by activated Wnt/PCP signalling. Abbreviations; Dvl – Dishevelled, Fzd – Frizzled, JNK – cJun N-terminal Kinase, Dkk1 – Dickkopf-1.

A recent report has suggested that A53T acts via JNK(55). This is not at odds with our hypothesis given the Wnt/PCP pathway bifurcates below Daam to signal via RhoA/ROCK to modulate cytoskeletal dynamics, and via JNK to modulate target gene transcription(56) (see schematic, figure 4).

Aβ acts on dendritic spines via Dkk1, which acts upon Wnt at the level of LRP6(34). Our observations indicate that oA53T-αSyn acts on Wnt/PCP at or above the level of *Daam1*, (schematic figure 4), raising the possibility that αSyn may also act at the level of LRP6, which is intriguing given the human αSyn gene, SNCA, is one of four autosomal dominant familial PD genes, and the other three, *LRRK2*, *GBA1*, and *VPS35*, have all been found to impact Wnt signalling precisely at the level of LRP6(57–59).

Further investigation is needed to clarify where exactly in the Wnt pathway A53T-αSyn acts to drive spine loss and to determine if Aβ and αSyn act in parallel through Wnt/PCP or in tandem. If the latter, it would indicate that the synaptic effects of Aβ might be αSyn-dependent, or the converse, either being of great importance for our better understanding of each of the neurodegenerative diseases in question here.

Of note, here we demonstrate that oA53T-αSyn acts on dendritic spines through the same pathway as Aβ and is blocked by fasudil. We are currently conducting a phase IIa clinical trial with fasudil for early-stage AD (ClinGov NCT06362707). Fasudil is off-patent and has a favourable safety profile making it a good candidate drug for repositioning(60). Several preclinical studies have previously found fasudil to be of benefit in rodent models of PD(61–64). Together, this indicates that fasudil may have benefit for treating PD and other synucleinopathies involving synapse loss, something others have previously suggested and are now testing (ClinGov NCT05931575)(65).

In summary, our observations indicate that Aβ and αSyn work in a common pathway to drive synapse loss. This gives credence to, and offers a molecular basis for, the many previous reports that have suggested a functional relationship exists between Aβ and αSyn. That Aβ and αSyn do appear to be functionally connected through Wnt/PCP suggests that Alzheimer’s and the synucleinopathies may share a common underlying disease mechanism - the dysregulation of Wnt/PCP signalling within dendritic spines. In addition to their impact on synapses, both Aβ and αSyn have detrimental effects on cerebrovascular function, affecting endothelia health and disruption of the blood-brain barrier (BBB). Given the importance of Wnt/β-catenin signalling in brain endothelial health and BBB maintenance(66, 67), it is possible Aβ and αSyn may also impact endothelia through disrupting canonical Wnt. This supports the growing view that rather than viewing PD, PDD, DLB and AD as separate disease entities they ought to be seen as different manifestations along a neurodegenerative disease continuum.

## Acknowledgements

This work was supported by the Rosetrees Trust and Race Against Dementia through award, RAD-2023-Full\1017, to Killick, Ribe, Hirth and Svenningsson; a Michael J Fox Foundation award, MJFF-025583, to Aarsland and Killick; and awards from the Swedish Parkinson Fund and the Swedish Brain Fund to Svenningsson.

